# TTYH family members form tetrameric complexes within the cell membrane

**DOI:** 10.1101/2021.09.05.459021

**Authors:** Emelia Melvin, Elia Shlush, Moshe Giladi, Yoni Haitin

## Abstract

The conserved Tweety homolog (TTYH) family consists of three paralogs in vertebrates, displaying a ubiquitous expression pattern. Although considered as ion channels for almost two decades, recent structural and functional analyses refuted this role. Intriguingly, while all paralogs, studied following detergent solubilization, shared a dimeric stoichiometry, their spatial organization differed. Here, we determined the stoichiometry of intact mouse TTYH (mTTYH) complexes in cells. Using cross-linking and single-molecule fluorescence microscopy, we demonstrated that mTTYH1 and mTTYH3 form tetramers at the plasma membrane. Blue-native PAGE and fluorescence-detection size-exclusion chromatography analyses revealed that detergent solubilization results in the dissolution of tetramers into dimers, suggesting a dimer-of-dimers assembly mode. As cross-linking analysis of the soluble extracellular domains also showed tetrameric stoichiometry, we explored the effect of membrane solubilization and disulfide bridges integrity and established their contribution to tetramer stability. Future studies of the native tetrameric TTYH characterized here may illuminate their long-sought cellular function.

## Introduction

The Tweety protein (TTY) was originally identified in the *drosophila melanogaster* flightless locus^1,2^. Subsequent studies identified three conserved TTY homologs (TTYH1-3) in vertebrates^3–5^, exhibiting a differential expression pattern. While TTYH3 is ubiquitously expressed^6^, the expression of TTYH1 and TTYH2 is limited to the central nervous system^7^ and testis^8^. While initial studies using drosophila knockout did not establish a clear functional role^3^, later studies suggested that this conserved family of membrane proteins serves as chloride channels, activated by either Ca^2+^ or cell swelling^6,9^. Specifically, it was suggested that TTYH1 mediates Cl^-^ currents in response to mechanical force^6,10^, while TTYH2 and TTYH3 isoforms underlie a Ca^2+^-dependent maxi-Cl^-^ current found in neurons^11^ and skeletal muscle^12^. Moreover, TTYH-mediated Cl^-^ currents were recorded using mammalian heterologous expression systems^6,13^.

Importantly, two recent studies elucidated the long-sought structure of TTYH family members^14,15^. However, strikingly, anion conductive pathways were not identified in any of the human (hTTYH) or mouse (mTTYH) members. Moreover, functional studies failed to reproduce the previously reported ion channel activity related to TTYH expression. The structures revealed a novel subunit topology common to all TTYH members, consisting of five transmembrane helices and a large a-helical extracellular domain (ECD). However, while the hTTYH paralogs and mTTYH3 were shown to exhibit a side-by-side dimeric organization involving interaction interfaces within the ECD and transmembrane regions, such an organization was shared by mTTYH2 only in the presence of calcium^14,15^. Conversely, it was suggested that in the absence of calcium, the dimers form by a head-to-head interaction between the ECDs of subunits from juxtaposing membranes^14^. Another surprising finding concerned the dimeric interface organization among the hTTYH paralogs, resulting from rigid body rotation along the two-fold axis of the individual monomers^15^.

Unfortunately, despite the extensive and detailed structural characterization of five different TTYH family members, their functional role remains debated^16^. Nonetheless, previous studies demonstrated that members of this family are involved in several pathophysiological processes^17^. For example, TTYH members were shown to dramatically change their expression pattern following Status Epilepticus^18^, as well as to drive colonization of gliomas^19^. Indeed, *ttyh1* was shown to be strongly upregulated in an array of childhood brain tumors^7^, with fusion of its promoter to the large microRNA cluster C19MC driving the development of a particularly aggressive form of pediatric cancer^20^.

As the structures of TTYH members, determined following detergent-mediated membrane solubilization, revealed distinct spatial organizations^14,15^, elucidation of their stoichiometry in the native membrane environment may hint towards the functional and cellular roles of TTYH members. This is of special importance given the involvement of TTYH members in human pathologies. Unexpectedly, using biochemical analyses, together with single molecule fluorescence microscopy^21^ and fluorescence-detection size-exclusion chromatography (FSEC)^22^, we show that TTYH family members form tetrameric complexes in the native membrane environment, which dissociate into dimers by detergent solubilization.

## Results

### TTYH members form high-order oligomeric assemblies in cells

To estimate the stoichiometry of TTYH proteins within their native membrane environment, we overexpressed FLAG-tagged mTTYH paralogs in HEK 293 cells (Fig. 1a). While we could clearly identify the expression of mTTYH1 and mTTYH3, a band corresponding to mTTYH2 could not be detected. Hence, we proceeded with investigation of mTTYH1 and mTTYH3 complex stoichiometry using *in* situ cross-linking with the amine-reactive, homobifunctional bis-sulfosuccinimidyl suberate (BS^3^)^23,24^. Importantly, BS^3^ is membrane impermeable, and thus can only cross-link primary amines of mTTYH subunits at the cell surface, exposed to the extracellular milieu. Incubation of the cells with BS^3^ resulted in the emergence of multiple migration species in both isoforms, with a mass culminating to that expected for tetrameric oligomers (∼240 kDa) (Fig. 1b, c). Together, these results suggest that mTTYH proteins may form higher-order oligomeric assemblies under native conditions and point towards a possible common tetrameric assembly for mTTYH1 and mTTYH3.

**Fig. 1.**
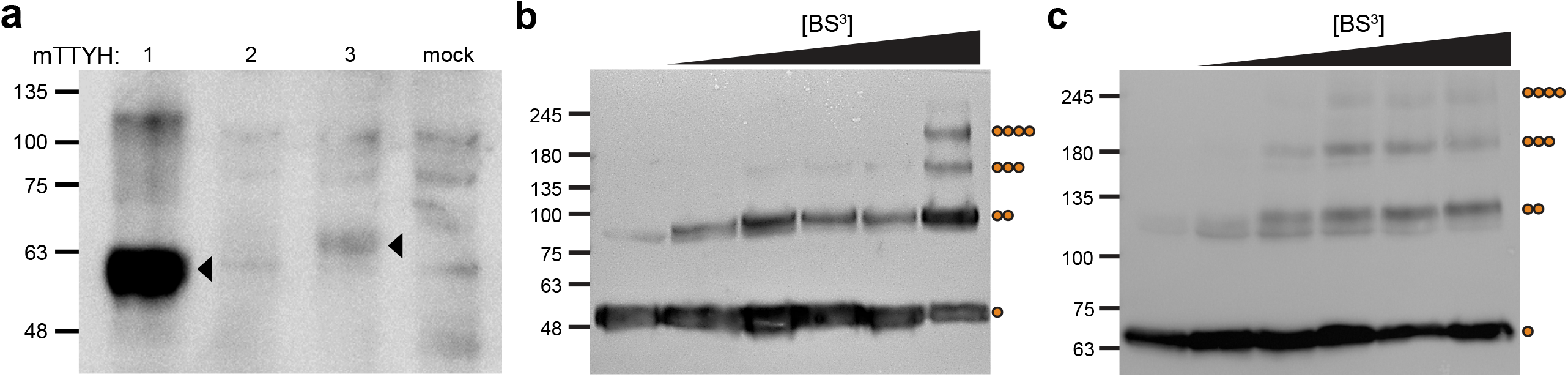
*In situ* cross-linking analyses of mTTYH family members. **a** SDS-PAGE western blot analysis of HEK 293 cells transiently transfected with the indicated FLAG-tagged mTTYH isoforms. Arrowheads indicate the bands corresponding to mTTYH1 and mTTYH3. Note that there is a nonspecific band in mTTYH2, which can also be found in non-transfected cells (mock). **b, c** SDS-PAGE western blot of mTTYH1 (**b**) and mTTYH3 (**c**) cross-linking with increasing concentrations of BS^3^. Molecular weights and oligomeric states are shown on the left and right, respectively.

### Tetrameric assembly of mTTYH complexes revealed by single-molecule subunit counting

While *in situ* cross-linking indicated that mTTYH proteins are composed of multiple subunits, we cannot exclude the possibility that heteromeric assembly formation of mTTYH dimers^14,15^ with additional endogenous proteins underlies the observed migration patterns. Thus, to directly examine the number of mTTYH subunits within a complex, in living cells, we resorted to the single-molecule subunit counting approach^21^. This method directly inspects the number of subunits within membrane-delimited complexes by monitoring the quantal bleaching of conjugated fluorophores. To this end, we generated C-terminal EGFP fusions of mTTYH1 and mTTYH3. Next, we titrated mTTYH-EGFP expression in *Xenopus* oocytes until a low enough level was achieved, enabling us to resolve individual spots using Total Internal Reflection Fluorescence Microscopy (TIRFM) (Fig. 2a, d). Cells were continuously illuminated, and time series images were recorded at 30 Hz until reaching baseline (100-200 seconds). Fluorescence intensities of stationary fluorescent spots were measured over the illumination duration, and bleaching steps were counted manually (Fig. 2b, e). As a positive control we used the EGFP fusion of KCNH1, a known tetrameric voltage-gated K^+^ channel^25^ (Fig. 2g, h, i). Importantly, both mTTYH1 and mTTYH3 exhibited bleaching profiles similar to that of KCNH1 (Fig. 2c, f, i), consistent with their tetrameric assembly. Indeed, the data were best fit by a binomial distribution^26^ assuming a 4-mer with a probability (*p*) of EGFP being fluorescent of 0.7 (Supplementary Fig. 1). Importantly, the stoichiometry we measured by subunit counting agrees well with the *in situ* cross-linking results.

**Fig. 2.**
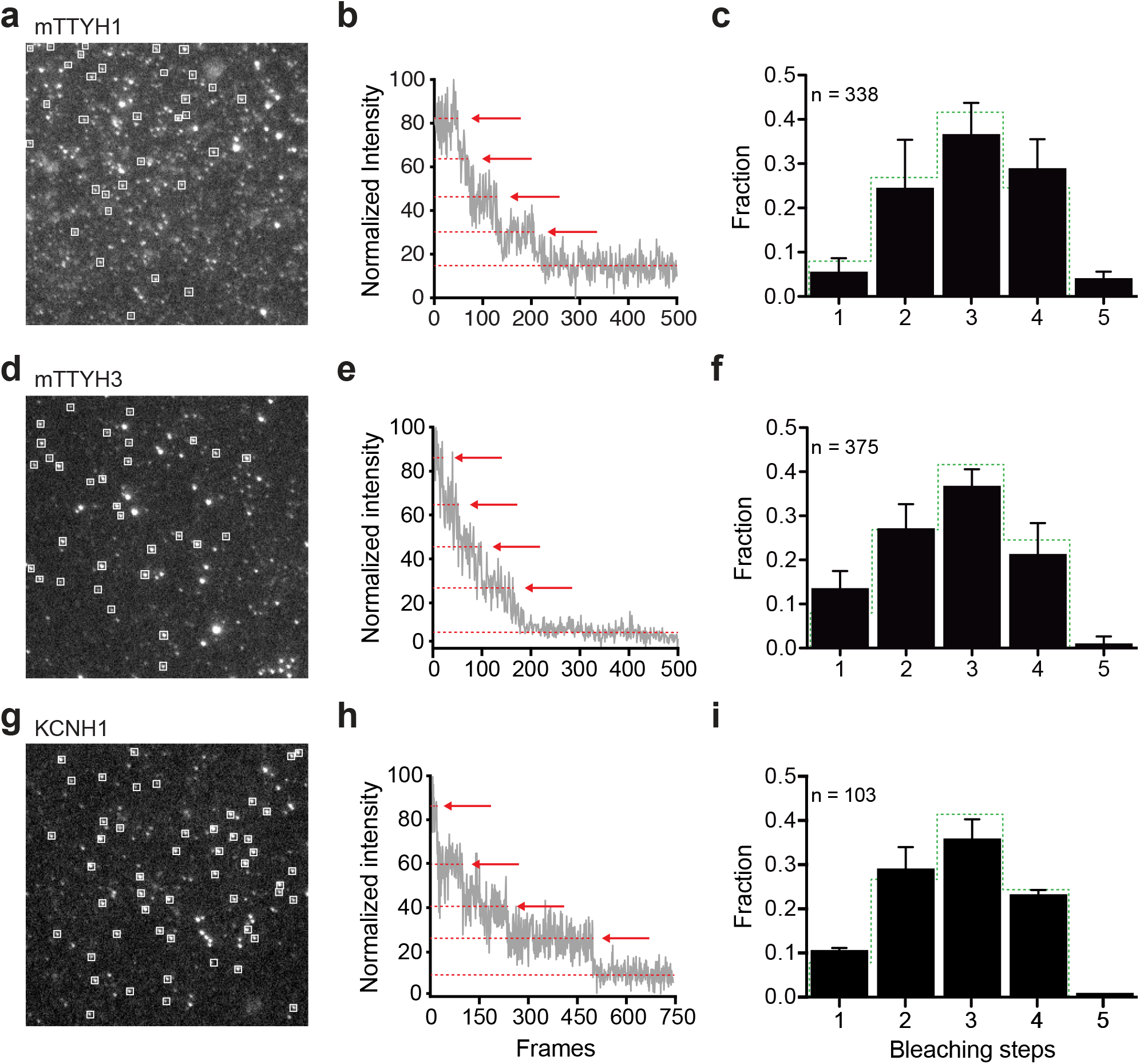
Single-molecule subunit counting of mTTYH family members. Analyses of EGFP-fused mTTYH1 (**a-c**), mTTYH3 (**d-f**), and KCNH1 (**g-i**). **a, d, g** Representative images sampled from 2 minutes movies, one day post injection of 1 ng cRNA, using TIRFM. Spots of interest are highlighted by white squares. **b, e, h** Representative fluorescence intensity traces obtained from single spots showing four steps of photobleaching, as indicated by arrows. **c, f, i** Probability distributions of bleaching steps (black bars), with error bars indicating SEM. Binomial distribution (*p* = 0.7) for tetrameric complexes is presented as dashed green line. n indicates the number of spots analyzed.

### Super resolution microscopy analyses of mTTYH complexes

Given the apparent discrepancy between the recent structural studies of TTYH members and the observed stoichiometry in the cellular context we sought to provide an independent assessment of mTTYH complexes sizes, as a surrogate for stoichiometry. Stochastic optical reconstruction microscopy (STORM) offers a sub-diffraction limit resolution imaging, enabling the visualization of individual complexes and assessment of their planar distribution and dimensions^27^. Specifically, STORM involves the repeated stochastic activation of single fluorescent molecules within a diffraction-limited spot, with the super-resolution image reconstructed from the calculated locations of each molecule^28^. Following fixation and permeabilization, cells expressing FLAG-tagged mTTYH1 and mTTYH3 were subjected to immunofluorescent labelling, followed by STORM (Fig. 3a, d). As expected, nearest-neighbor distances analysis^27^, reflecting the distance between individual fluorophores, demonstrated a distribution with a sharp peak at 10–20 nm, the theoretical resolution limit of STORM, for both mTTYH1 and mTTYH3 (Fig. 3a, e). Specifically, mTTYH1 and mTTYH3 showed peak distributions at 9.11 ± 0.01 and 8.76 ± 0.02 nm, with peak amplitudes of 27.58 ± 0.03% and 28.12 ± 0.08%, respectively. Next, to derive complexes sizes from the nearest neighbor distance analysis, we performed a spatial clustering of localizations analysis^29^ (Fig. 3c, f). This analysis revealed a prominent clustering distribution for both mTTYH1 and mTTYH3, with a peak at a radius corresponding to that expected from the theoretical arrangement of a multimeric protein-antibodies complex^27,29^. Specifically, mTTYH1 and mTTYH3 demonstrated mean cluster size of 41.64 and 50.46 nm, respectively. Notably, similar radii were measured before using STORM for tetrameric KCNQ channels^27^. Therefore, STORM data further support the tetrameric stoichiometry of mTTYH protein in cells.

**Fig. 3.**
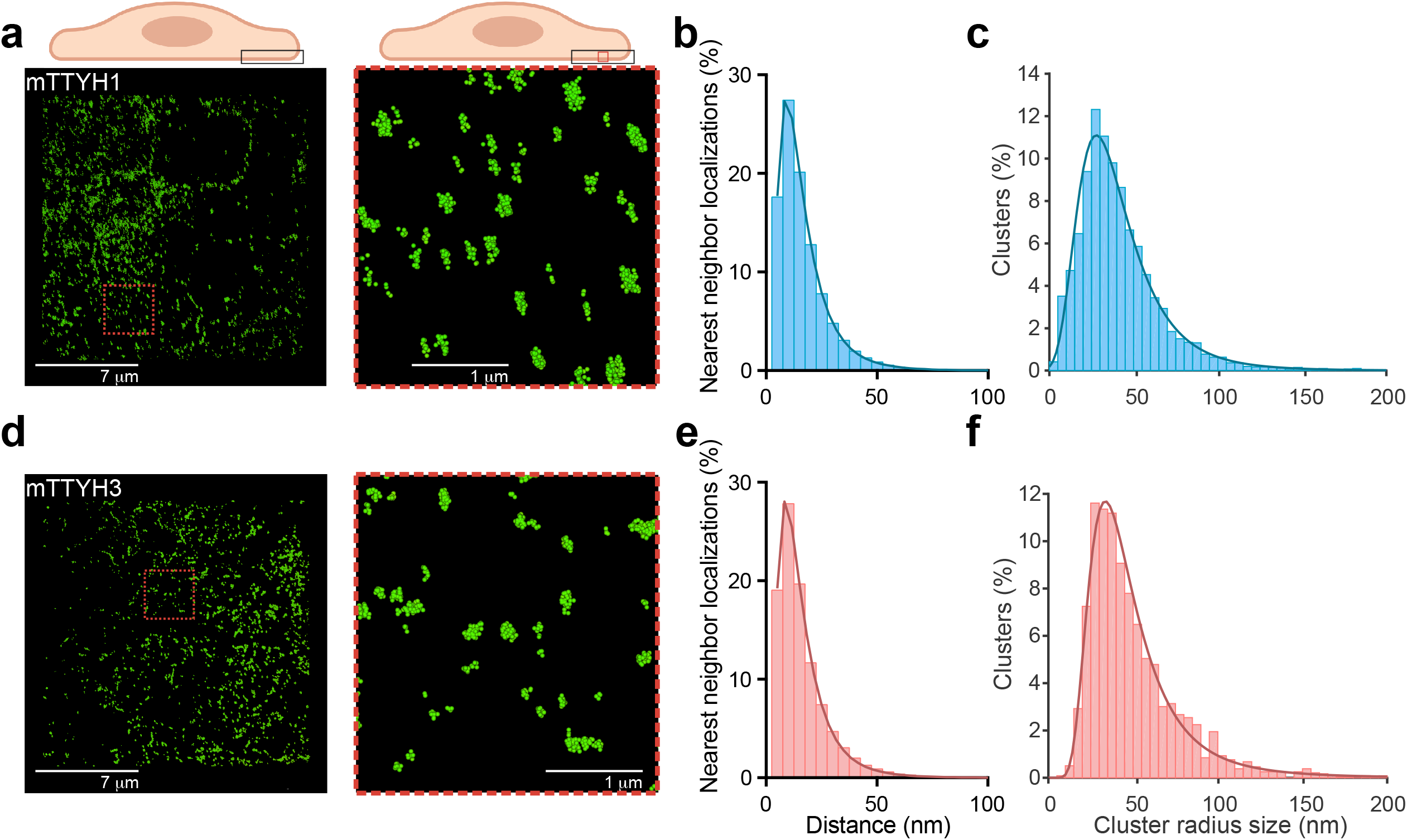
mTTYH complexes observed by STORM. **a, d** STORM imaging of HEK 293 cells transiently transfected with FLAG-tagged mTTYH1 (**a**) or mTTYH3 (**d**). Representative Z-axis truncated datasets (left) and blowout images (right) are provided. **b, e** Nearest neighbor distances distribution for cells expressing mTTYH1 (**b**) and mTTYH3 (**e**). Curves were fit using Log (Gaussian) nonlinear equation. **c, f** DBSCAN algorithm cluster radii probability of mTTYH1 (**c**) and mTTYH3 (**f**). Data were fit by generalized extreme value distribution.

### Detergent solubilization of mTTYH complexes results in the emergence of a dimeric population

In contrast to the tetrameric stoichiometry, we observed in the cellular environment, the recent structures of members of the TTYH family shared a dimeric organization. However, as these structures were determined following membrane solubilization by detergents, we hypothesized that this process may result in the dissociation of TTYH complexes. Therefore, we used Blue Native PAGE (BN-PAGE) to assess subunit stoichiometry following detergent solubilization using n-Dodecyl-β-D-Maltopyranoside (DDM) (Fig. 4a). This analysis revealed that both mTTYH1 and mTTYH3 display a complex migration pattern, consisting of dimeric (∼120 kDa) and tetrameric (∼240 kDa) populations (Fig. 4a), indicating detergent-mediated perturbation of the quaternary structure of the complex. Conversely, the dimeric population may arise from the reagents used for the BN-PAGE analysis (e.g. Coomassie stain). Thus, to examine whether the dimeric population stems from a detergent-mediated tetramers disassembly, we used FSEC. HEK 293 cells expressing EGFP fusions of mTTYH1 or mTTYH3 were solubilized with DDM and subjected to FSEC analyses (Fig. 4b). Intriguingly, poly-dispersed elution profiles were evident for both isoforms, with peaks detected at volumes corresponding to tetrameric as well as dimeric assemblies. Specifically, mTTYH1 displayed an elution profile mainly corresponding to that expected for dimers, while mTTYH3 dispersion was skewed towards an elution volume expected for tetramers (Fig. 4b).

**Fig. 4.**
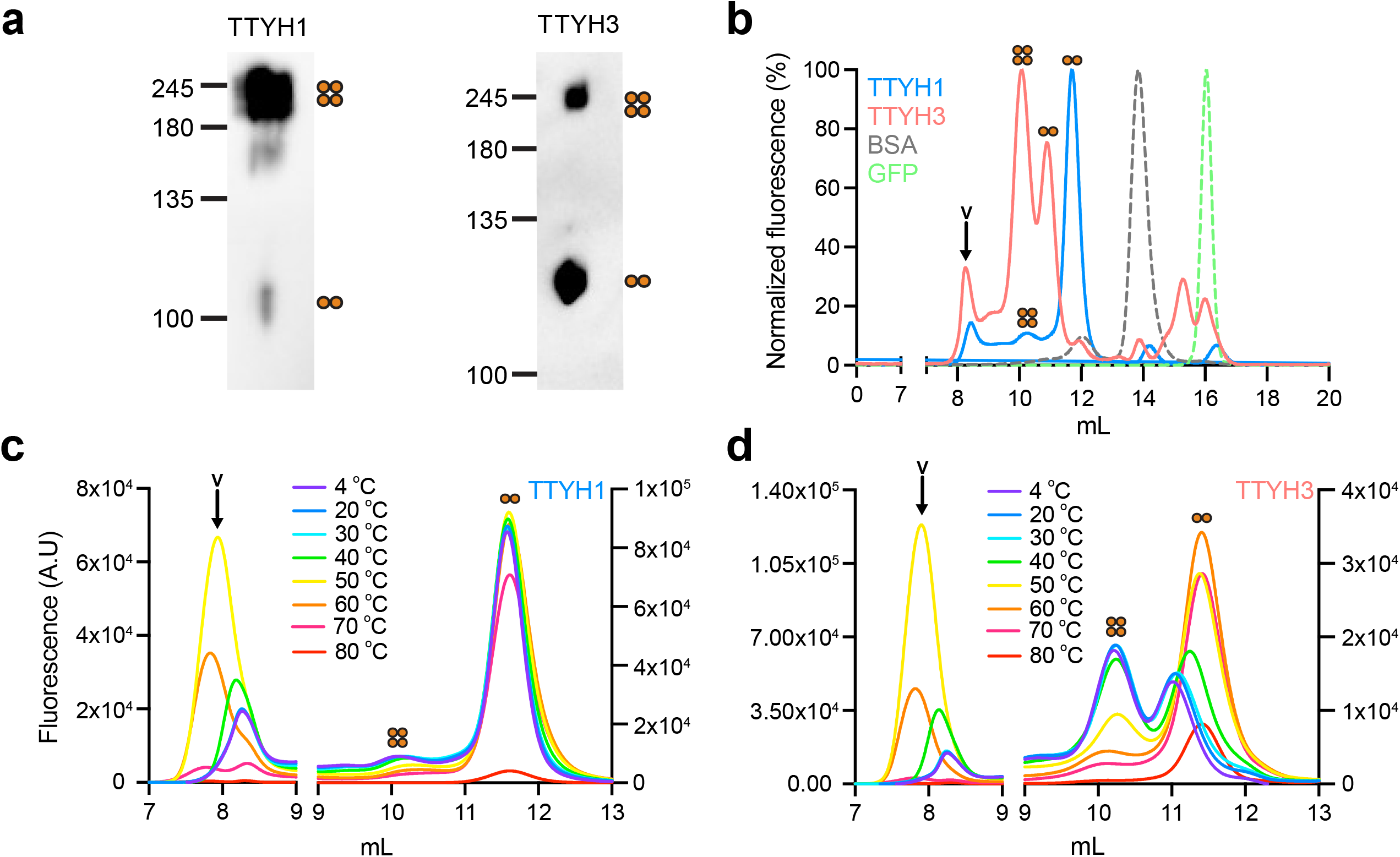
Analyses of detergent-solubilized mTTYH isoforms. **a** BN-PAGE western blot analysis of HEK 293 cells transiently transfected with mTTYH1 (left) or mTTYH3 (right). **b** FSEC chromatograms depicting the elution profiles of DDM solubilized HEK 293 cells expressing the indicated EGFP-fused mTTYH isoforms. BSA and EGFP profiles are provided for reference. **c, d** FSEC chromatograms of mTTYH1 (**c**) and mTTYH3 (**d**) pre-incubated for 10 minutes at the indicated temperatures.

To gain further insights into the dissociation of mTTYH tetramers, we studied their thermal denaturation using FSEC-based thermostability assay (FSEC-TS)^30^. Samples were subjected to 10-minute thermal insults, ranging from 20 °C to 80 °C, followed by centrifugation and FSEC analysis (Fig. 4c, d). With increasing temperatures, the population distribution shifted towards dimers (Fig. 4c, d, Supplementary Fig. 2), accompanied by the emergence of soluble aggregates eluting at the column void volume. These results suggest that mTTYH complexes may feature a dimer-of-dimers assembly, with a detergent-resistant dimeric interface and a labile surface involved in tetramerization.

### The tetrameric interface of mTTYH is stabilized by cholesterol and intra-subunit disulfides

The possibility that mTTYH complexes arise from a dimer-of-dimers assembly led us to examine whether the tetramer/dimer ratio can be modified by the solubilization conditions. Thus, we solubilized the cells with DDM in the presence a of cholesteryl hemisuccinate (CHS). CHS is known to improve detergent extraction efficiency and monodispersity in structural studies of membrane proteins^31^. Indeed, the inclusion of CHS in the solubilization buffer increased the tetramer/dimer ratio, as determined by FSEC, for both mTTYH1 and mTTYH3 (Fig. 5). This suggests that CHS can stabilizes the native cell-membrane residing tetramers.

**Fig. 5.**
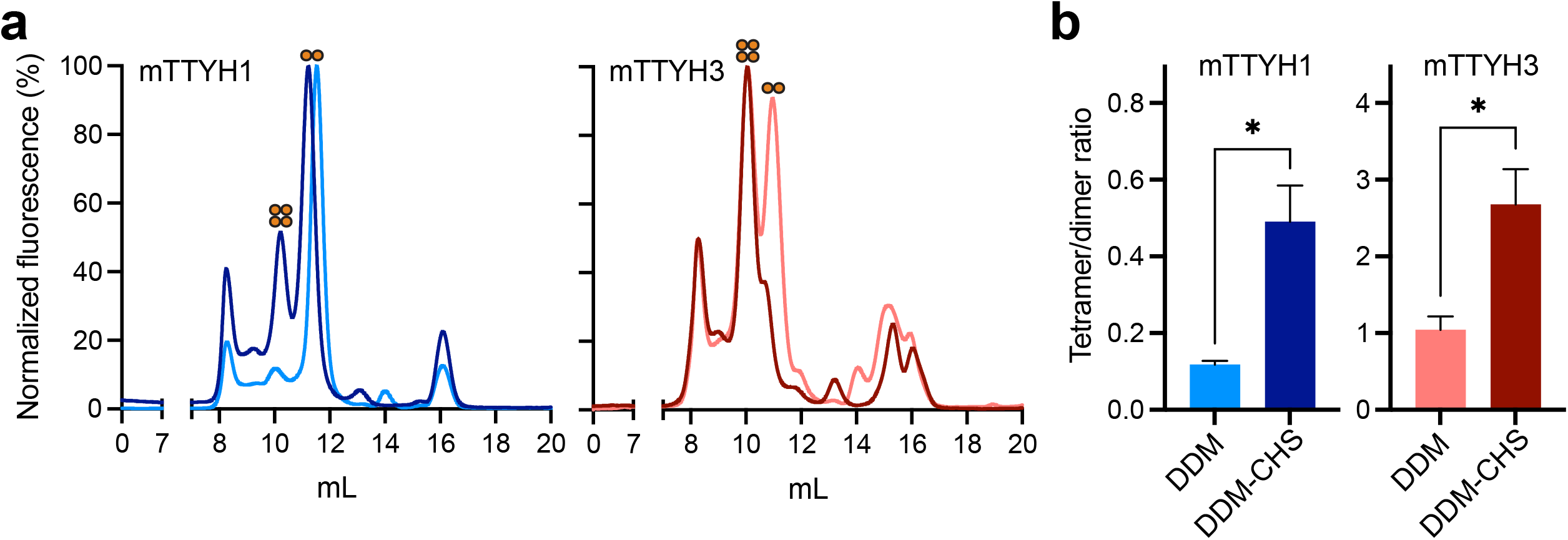
Solubilization by DDM/CHS stabilizes mTTYH tetramers. **a** FSEC chromatograms depicting the elution profiles of DDM/CHS solubilized HEK 293 cells expressing the indicated EGFP-fused mTTYH paralogs. **b** Tetramer/dimer ratio of the indicated mTTYH paralogs following solubilization with DDM or DDM/CHS (n=3, * *p* < 0.05).

Next, as solubilization with the DDM/CHS mixture increases the tetrameric fraction in the population of both mTTYH paralogs, we sought to determine if additional conditions may drive the dissociation of tetramers into dimers. The recent structure of both hTTYH and mTTYH paralogs revealed large interaction interfaces between the ECDs. Moreover, the structures show that the ECD contains two disulfide bridges, formed between conserved cysteine residues (Fig. 6). These bridges stabilize a helical flap-like hairpin fold. We reasoned that perturbation of these conserved structural elements may destabilize the ECDs and thereby the interaction interfaces between the subunits. Therefore, following solubilization with DDM/CHS, FSEC was performed before and after incubation with 10 mM 2-mercaptoethanol (2-ME) (Fig. 6). Importantly, 2-ME resulted in a marked shift towards dimers, further supporting the notion that the dimers arise from dissociation of tetramers, highlighting the role of the ECDs in complex assembly.

**Fig. 6.**
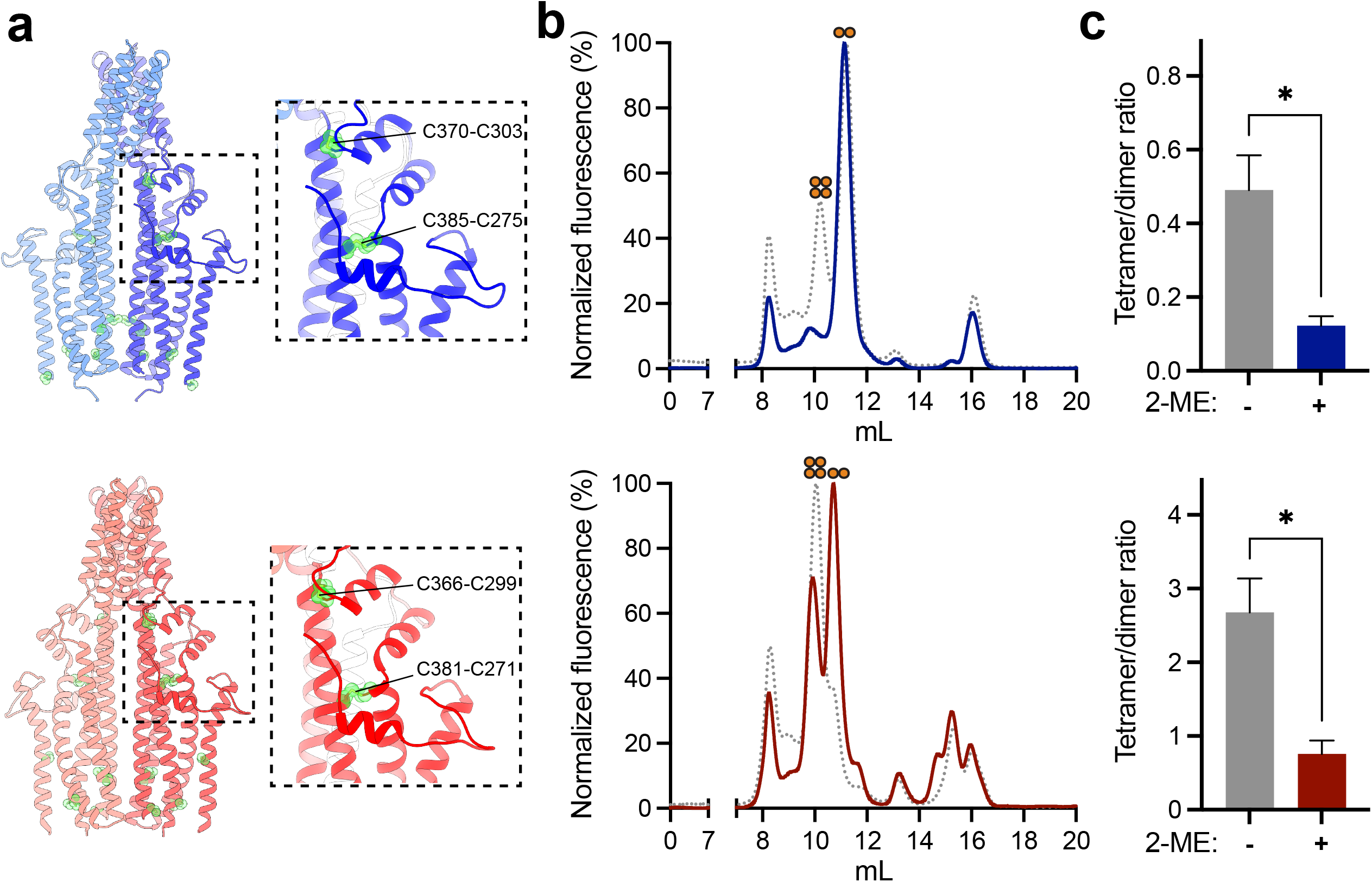
mTTYH disulfides contribute to tetramerization. **a** Cryo-EM structures of hTTYH1 (upper panel; PDB: 7P5J) and hTTYH3 (lower panel; PDB: 7P5C) in cartoon representation. The dashed rectangles frame zoom perspectives of the disulfide bridges within the ECDs. Cysteine residues are colored green and shown as spheres. **b** FSEC chromatograms depicting the elution profiles of DDM/CHS solubilized HEK 293 cells expressing the indicated EGFP-fused mTTYH1 (upper panel) and mTTYH3 (lower panel) before (gray curve) and following incubation with 10 mM 2-ME (colored curve). **c** Tetramer/dimer ratio of mTTYH1 (upper panel) and mTTYH3 (lower panel) before and following incubation with 2-ME (n=3, * *p* < 0.05).

### Isolated mTTYH3 ECDs form tetramers in solution

The structures of the intact TTYH proteins revealed disulfide bridges within the extracellular domains, which appear to contribute to tetramerization based on the effect of 2-ME (Fig. 6). Thus, to establish the role of the ECDs in tetramerization, we designed a construct consisting of the isolated domains (Fig. 7a). Specifically, focusing on mTTYH3, we introduced a T4 lysozyme (T4L) sequence between ECD1 (positions 107-209) and ECD2 (positions 260-386) to increase protein stability and solubility^32^. Due to the presence of disulfide bridges and glycosyl moieties^14,15,33^, we expressed this FLAG-tagged construct as a secreted protein in insect cells. Indeed, western blot analysis of the cell lysate revealed a complex migration pattern consistent with different stages in the protein biogenesis pathway. Conversely, a single band was detected in the secreted fraction obtained from the clarified lysate (Fig. 7b, left panel). Indeed, the secreted fraction is glycosylated, as determined by the change in the band mobility following treatment with Peptide:N-glycosidase (PNGase) F (Fig. 7b, right panel). The secreted protein was subjected to cross-linking analysis using the homo-bifunctional amine reactive cross-linker disuccinimidyl suberate (DSS), resulting in the emergence of higher-order oligomers, culminating to the size expected for a tetramer (Fig. 7c). Thus, together with the results obtained using intact TTYH (Fig. 6), we suggest that the ECDs participate in TTYH tetramerization.

**Fig. 7.**
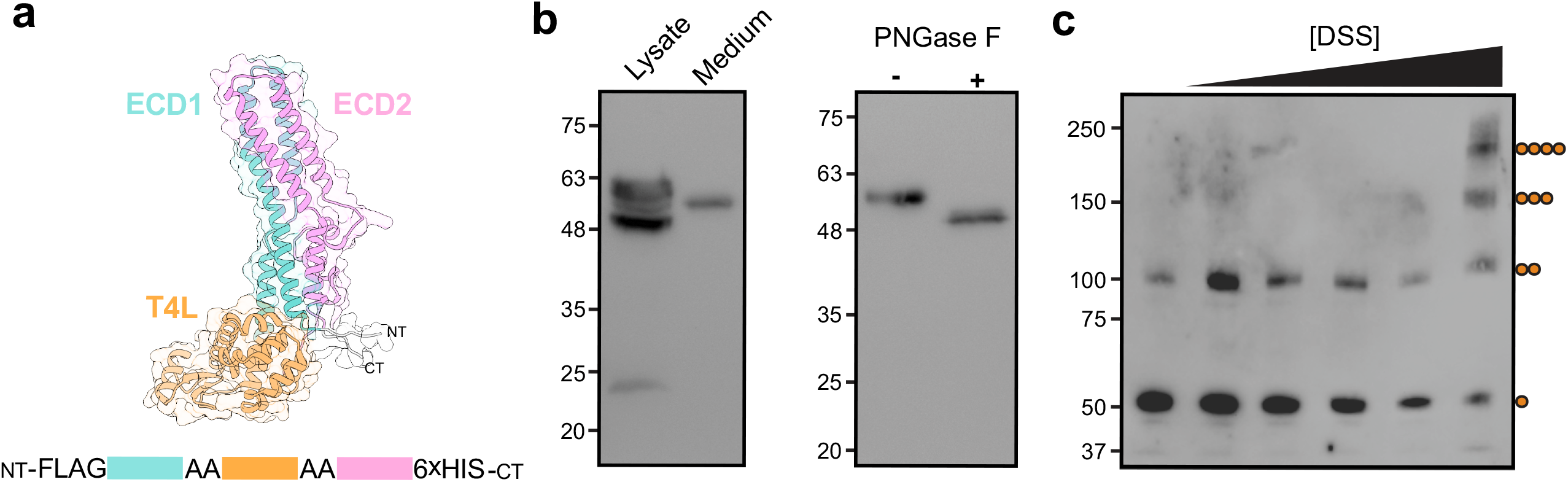
mTTYH3 ECDs form tetramers in solution. **a** A model of ECD1-T4L-ECD2 generated using AlphaFold2^44^. **b** Western blot analyses of mTTYH3 ECD1-T4L-ECD2 expression using anti-FLAG. Note the single band observed in the secreted fraction (Medium), compared with the multiple intracellular forms (Lysate) (left panel), and the effect PNGase F treatment (500 units) (right panel). **c** Cross-linking using increasing DSS concentrations (0 - 2000 µM) resulted in the exposure of higher oligomeric assemblies, culminating in a mass corresponding to that expected for a tetramer.

## Discussion

Over the past years the TTYH family gained attention mainly due to its expression in the CNS^7,8,34^ and association with various pathological conditions^18–20,34^. These observation motivated functional and structural studies^17^, culminating in the recent structural elucidation of hTTYH and mTTYH members^14,15^. However, while these structures lack an obvious Cl^-^ conductive pathway, contradicting the originally assigned functional role of these proteins as ion channels, the functional role of the TTYH family remains enigmatic. Here, we performed a comprehensive analysis of mTTYH stoichiometry in their natural settings. Our data suggest that TTYH channels exist in membrane as tetramers, and expose the impact of detergent solubilization on complex integrity.

To demonstrate that mTTYH form multi-subunit complexes, we used *in situ* cross-linking. This analysis revealed mTTYH oligomers at the plasma membrane, corresponding to tetrameric assemblies (Fig. 1b, c). However, many membrane protein complexes exhibit a heteromeric assembly with auxiliary subunits^35–38^. Indeed, it was previously shown that members of the TTYH family can undergo ubiquitination^39^, possibly contributing to the observed mass. Therefore, it remained possible that the high molecular weight bands observed following the BS^3^ *in situ* cross-linking analyses harbor additional proteins, accounting for the observed mass, rather than representing mTTYH homotetramers. Thus, to directly assess the number of subunits within mature complexes, in their undisrupted environment, we resorted to the use of single-molecule fluorescence microscopy approaches.

Consistent with the *in situ* cross-linking analysis, two complementary approaches independently support the tetrameric nature of mTTYH complexes. Indeed, measurements of the photobleaching steps distribution in subunit counting experiments^21^, for both isoforms, was in striking agreement with that measured for KCNH1 (Fig. 2), a K^+^ channel with well-established tetrameric stoichiometry^25^. Moreover, to assess the complex dimensions at the plasma membrane, serving as a surrogate for its stoichiometry, we proceeded with STORM analysis. Comparison with previous STORM microscopy studies of the tetrameric KCNQ voltage gated K^+^ channel^27^ revealed nearly identical cluster size and distribution compared with those of mTTYH1 and mTTYH3, reported here (Fig. 3). As the molecular weights of KCNQ and mTTYH subunits are similar, it can be deduced that mTTYH complexes share a similar tetrameric stoichiometry. Together, our multi-faceted investigations unequivocally argue that four mTTYH subunits assemble to form the native complexes and that the high molecular weight bands observed using the *in situ* cross-linking analysis indeed represent mTTYH homotetramers.

The recent cryo-EM structures of TTYH family members revealed a common dimeric stoichiometry, in sharp contrast to the tetrameric assemblies observed here, in the cellular milieu. Puzzled by this apparent discrepancy, we sought to determine the key underlying difference between the experimental systems. Intriguingly, previous BN-PAGE and FSEC analyses of the bona-fide tetrameric GluA1 and GluA2 revealed that, following detergent-solubilization, tetrameric and dimeric populations coexist^40^, in accordance with their dimer of dimers assembly mode^41^. Additionally, a study of the human excitatory amino acid transporter (EAAT) revealed that detergent solubilization significantly destabilizes the inter-subunit interface^42^. Indeed, BN-PAGE analyses of detergent-solubilized mTTYH complexes expose that, additional to the tetrameric population, a significant dimeric fraction was evident (Fig. 4a). To further explore the tetramer to dimer dissociation, we performed FSEC analysis of EGFP-fused mTTYH subunits, following solubilization with DDM. Indeed, the elution profile of mTTYH1 mainly consisted of dimers and that of mTTYH3 displayed roughly 3:2 ratio of tetramers to dimers (Fig. 4b). Moreover, to determine whether tetramers can dissociate into dimers, we explored the thermal stability of the detergent-solubilized mTTYH complexes population using FSEC-TS. We observed a temperature-related shift in the stoichiometry of both mTTYH1 and mTTYH3. With an increased insult temperature, the dimeric population amplitude increases on the expense of the tetrameric one, followed by a decrease in the fluorescence amplitude of the dimers at high temperatures. Importantly, we could not detect the emergence of a putative monomeric population at any insult temperature for either mTTYH1 or mTTYH3. These results suggest that the tetramers dissociate into dimers, which are the obligatory indivisible building block of the mTTYH complexes, as observed in the cryo-EM structures^14,15^.

Following the demonstration of mTTYH tetramer destabilization by detergent solubilization and temperature insults, we tested if the tetramers could be stabilized during solubilization. To that end, we using a DDM/CHS mixture, as CHS is known for its stabilizing effect in membrane protein purification^31^. The inclusion of CHS in the solubilization step indeed increased the tetramer/dimer ratio (Fig. 5), further supporting the notion that dimers arise from the tetramer dissociation. Finally, we wondered if the tetramer is stabilized by the interaction between the ECDs, as observed in the cryo-EM structures of the dimers. These structures revealed that the ECDs are stabilized by two intra-subunit disulfide bridges. Hence, we examined the tetramer/dimer ratio following the incubation with the reducing agent 2-ME, which resulted in a decreased ratio. This observation supports the importance of intact disulfides for the tetramerization surfaces of mTTYH complexes. Indeed, cross-linking of isolated ECDs unveiled the formation of higher-order oligomers (Fig. 7c), consistent with tetramers, further supporting the stoichiometry observed in cellular membranes, and underscoring the role of these soluble domains in complex formation.

In conclusion, here we expose for the first time that mTTYH form homo-tetramers at the plasma membrane using *in situ* cross-linking and single molecule fluorescence microscopy. Our analysis suggests a dimer of dimers assembly mode, which involves the ECDs. Moreover, this tetrameric assembly is detergent-sensitive and is affected by the solubilization conditions. Importantly, our findings should guide future structural investigations of TTYH structure in the tetrameric context, which may illuminate the function of these enigmatic proteins.

## Acknowledgments

This work was performed in partial fulfillment of the requirements for a Ph.D. degree of E.M., Sackler Faculty of Medicine, Tel-Aviv University, Israel. This work was supported by the Israel Science Foundation (grant 1721/16) (Y.H.), the Israel Cancer Research Fund grants 01214 (Y.H.) and 19202 (M.G.), and from the German-Israeli Foundation for Scientific Research and Development (grant No. I-2425-418.13/2016) (Y.H.). Support also came from the I-CORE Program of the Planning and Budgeting Committee and the Israel Science Foundation (grant 1775/12) (Y.H.), Recanati Foundation (M.G.), Marguerite Stolz Research Fellowship (Y.H.), Kahn foundation’s Orion project, Tel Aviv Medical Center, Israel (M.G.), Israel Cancer Association (grant 20200037) (Y.H. and M.G.), and the Claire and Amedee Maratier Institute for the Study of Blindness and Visual Disorders, Sackler Faculty of Medicine, Tel-Aviv University (Y.H. and M.G.).

## Author Contributions

Conceptualization, M.G. and Y.H.; Methodology, M.G. and Y.H.; Investigation, E.M., E.S.; Formal Analysis, E.M., E.S., M.G., Y.H.; Writing – Original Draft, E.M., M.G., Y.H.; Writing – Review & Editing, E.M., M.G., Y.H.; Supervision, M.G., Y.H.; Funding Acquisition, M.G., Y.H.

## Methods

### DNA Constructs and Expression

For FSEC, mTTYH1 and mTTYH3 were subcloned into the pEGFP-N1 multiple cloning site (MCS) using the Gibson assembly method^43^. For BN-PAGE, cross-linking, and STORM experiments, the same constructs were modified with the introduction of a FLAG tag instead of the original EGFP sequence. For subunit counting experiments, the intact mTTYH-EGFP fusions were cloned into the MCS of the pGH19 vector. For expression of the isolated mTTYH3 ECDs, the ECD1-T4L-ECD2 construct (Fig. 7a) was subcloned seamlessly between an N- and C-terminal FLAG and hexa-histidine tags, respectively, into the pK503-9 plasmid.

### Cross-linking analysis

HEK 293 cells were transiently transfected with FLAG-tagged mTTYH constructs using calcium phosphate precipitation. 36 hours post-transfection, cells were washed twice with PBS, followed by cross-linking of the adherent cells by incubation with BS^3^ (Thermo Fisher Scientific) at 10-2000 μM for one hour at room temperature (RT). Subsequently, the reactions were quenched by the addition of Tris-HCl (pH 7.5) at a final concentration of 20 mM for 15 minutes. Cells were collected and centrifuged at 500 x g for 3 minutes, resuspended with lysis buffer (150 mM NaCl, 50 mM HEPES, pH 7.5, 1% (w/v) DDM (Anatrace), 1 mM PMSF), incubated for 1 hour at 4 °C and clarified by centrifugation at 21,000 x g for 10 minutes. Finally, the supernatant was subjected to western blot analyses using mouse anti-FLAG (M2) (Sigma).

### Single-molecule subunit counting

cRNA were synthesized *in vitro* from linearized DNA of EGFP-fused mTTYH constructs using HiScribe T7 ARCA mRNA Kit (New England Biolabs). 1–2 ng of cRNA were injected in *Xenopus laevis* oocytes and imaged after 18 hours of incubation at 18 °C. Imaging was performed using a Total Internal Reflection Fluorescence Microscopy (TIRFM) system (Zeiss)^21^. Briefly, prior to the experiment, cells were manually devitellinized and placed on high refractive index coverslips with the animal pole in front of a Zeiss alpha Plan-Fluar 100X / 1.45 NA oil immersion objective, at RT. EGFP-fused mTTYH subunits were excited using an argon 488 (100 mW) laser and fluorescence was obtained using 500 nm long-pass dichroic mirror in combination with a 525/50 nm band-pass filter. Images were collected at 30 Hz using cooled electron-multiplying charge-coupled device camera (Photometrics). Only isolated and immobile diffraction-limited spots were subjected for bleaching steps analysis. Fluorescence intensity of each spot was plotted over time and the bleaching steps were manually counted. 338, 375, and 100 spots were counted, from 2-6 oocytes using three different batches, for mTTYH1, mTTYH3 and KCNH1.

### Immunofluorescence staining for STORM

For imaging, acid washed, and poly L-lysine-coated 25 mm circular coverslips were used. HEK 293 cells were seeded (∼10^5^ cells/well) and transiently transfected the next day with FLAG-tagged mTTYH constructs, followed by incubation for 36 hours. Next, cells were washed with PBS and fixed with 4% (w/v) paraformaldehyde for 20 minutes at RT, permeabilized using 0.2% (w/v) triton X-100 for 5 minutes, and incubated in blocking buffer (5% (v/v) goat serum, 0.1% (w/v) BSA and 0.2% (w/v) triton X-100 in PBS) for one hour at RT, followed by incubation with mouse anti-FLAG (M2) (Sigma). Finally, the primary antibody was washed, cells were incubated with an anti-mouse Alexa Fluor 647 conjugated secondary antibody, and coverslips were stored in PBS at 4 °C protected from light until use.

### STORM imaging and analysis

Images were acquired using a Vutara 352 super resolution system, equipped with a 60X / 1.20 NA water immersion objective. Coverslips were imaged using an Attofluor cell chamber stage adapter (Invitrogen), after being layered with freshly prepared imaging buffer (50 mM Tris-HCl, pH 8.0, 10 mM NaCl, 10% (w/v) glucose, 20 mM cysteamine, 10% v/v β-mercaptoethanol, 1688 μg/ml glucose oxidase, and 140 μg/ml catalase). The 647 nm excitation laser line was used at 40% transmission output. 10,000 frames were recorded per a region of interest, following by analysis using the Vutara SRX 6.02.02 software. To include only intact complexes, localized at or in the vicinity of the plasma membrane, the Z-axis was truncated to 100 nm depth using the pre-determined calibration values obtained by prior measurements of Alexa Fluor 647 fluorescent beads. Next, nearest neighbor analysis with maximal radius of 500 nm was performed in cells expressing mTTYH1 or mTTYH3, and the distances containing 95% of the data was determined. These distances were used for as inputs for the DBSCAN algorithm^29^. DBSCAN clustered datasets were generated for all cells analyzed. Radii sizes of clusters were calculated, and data histograms were fit by generalized extreme value distribution.

### BN-PAGE

HEK 293 cells were transiently transfected with FLAG-tagged mTTYH constructs using the calcium phosphate precipitation method. 48 hours post transfection, cells were washed twice in phosphate buffered saline (PBS) and lysed with a buffer containing 50mM NaCl, 20 mM Tris-HCl, pH=7.5, 0.05% (for mTTYH1) or 0.1% (for mTTYH3) (w/v) DDM (Anatrace), protease inhibitor cocktail set III (Merck Millipore), 1mM phenylmethylsulfonyl fluoride (PMSF). Cell lysates were cleared by centrifugation (21,000 x g for 30 min, 4°C). Following determination of total protein concentration, equal quantities of protein were diluted 3:1 with native-PAGE sample buffer (50 mM Tris-Acetate, pH=7, 6 M HCl, 50 mM NaCl, 10% (w/v) glycerol, 0.001% (w/v) ponceau S and 0.25% (w/v) Coomassie G), and were analyzed using 10% Tris-Acetate gels and western blot with mouse anti-FLAG (M2) (Sigma).

### FSEC and FSEC-TS analyses

HEK 293 cells were transiently transfected with the mTTYH-EGFP fusion constructs. After incubation for 48 hours, cells were harvested by gentle pipetting, washed with PBS, and resuspended in 500 μl solubilization buffer (150 mM NaCl, 50 mM Tris-HCl, pH 7.5, 1% (w/v) DDM (Anatrace), protease inhibitor cocktail set III (Merck Millipore), 1 mM PMSF). When indicated, CHS (Anatrace) was added at 0.1% (w/v). Resuspended cells were rotated for 1 hour at 4 °C, followed by centrifugation at 21,000 x g for 1 hour. 50 μl of the supernatant were loaded onto a Superdex 200 Increase 10/300 GL column (Cytiva) pre-equilibrated with size exclusion buffer (150 mM NaCl, 50 mM Tris-HCl, pH 7.5, 1 mM DDM (Anatrace)) in line with a fluorescence detector (RF-20A; Shimadzu Scientific Instruments), with λexcitation / λemission of 488 / 507 nm, respectively. For FSEC-TS, clarified samples were incubated at the indicated temperatures for 10 minutes using a thermal cycler, and then centrifuged at 21,000 x g for 10 minutes, prior to the FSEC analysis described above.

### Isolated TTYH3 ECD expression in insect cells

Recombinant baculoviruses for expression of the mTTYH3 ECD1-T4L-ECD2 construct were generated using the Bac-to-Bac system (ThermoFisher Scientific). Sf9 cells were cultured at 27 °C in 1 L culture flasks with ESF-921 insect cell culture medium (Expression systems). Next, cells were infected at 1×10^5^ cells/ml with a viral ratio of 1:5 and incubated 72 hours post-infection. The media, containing the secreted protein, was clarified by centrifugation at 1200 × g for 5 minutes.

