## Supplementary information for "TTYH family members form tetrameric complexes within the cell membrane"

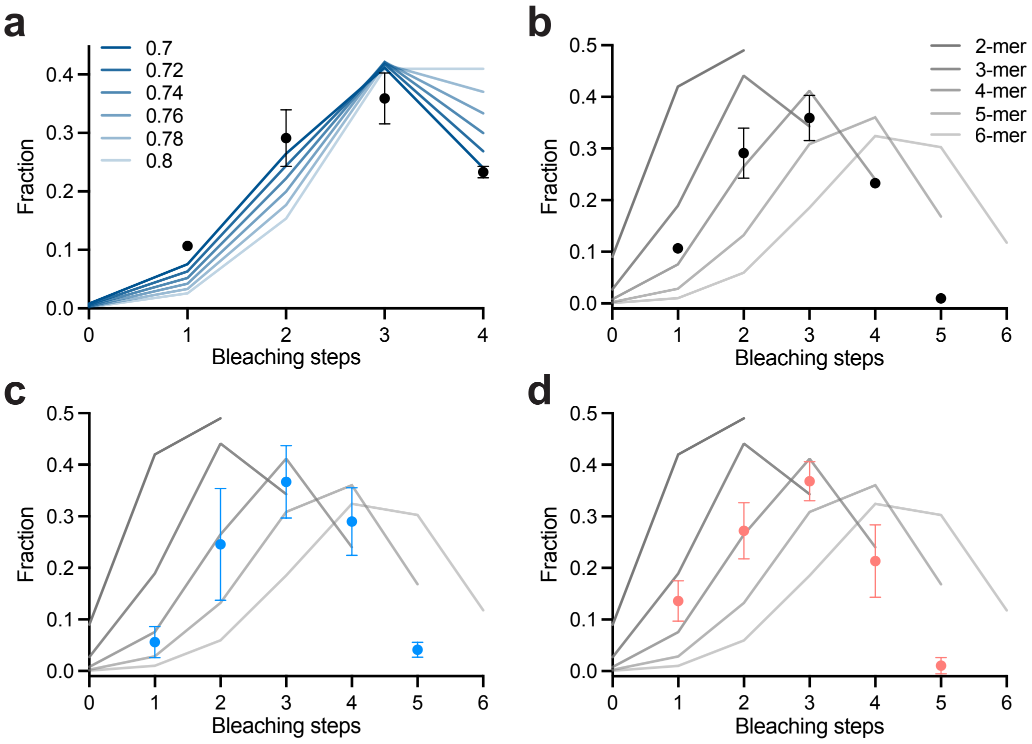


**Supplementary Fig. 1. Subunit counting data binomial fit. a** Determination of the EGFP probability of being fluorescent (*p*). Data collected from oocytes expressing homotetrameric KCNH1 channels was fitted with the expected distribution of tetrameric channels containing 1, 2, 3 or 4 functional GFP tags, allowing the probability that the EGFP tag is fluorescent to be a free parameter. A probability of 0.7 of the EGFP to be fluorescent matches well the observed data. **b-d** Distribution of the number of bleaching steps observed from oocytes expressing KCNH1 (**b**), mTTYH1 (**c**), and mTTYH3 (**d**). Circles, the average percentage of spots that bleached in each number of bleaching steps. Binomial equation fits, assuming the number of monomers in the complex and with the percentage of fluorescent EGFP molecules being a free parameter (graded grey shaded lines).

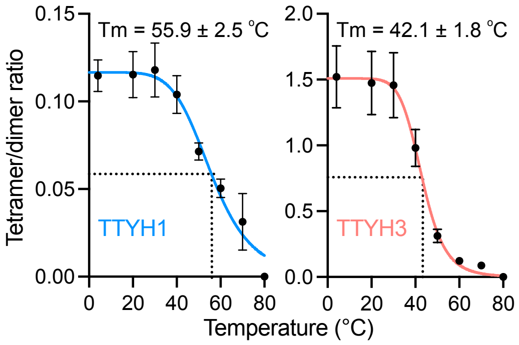


**Supplementary Fig. 2. Thermal denaturation of mTTYH paralogs.** Melting curves of mTTYH1 (left) and mTTYH3 (right). Melting temperatures (Tm) for tetramer/dimer ratio were calculated by fitting the curves with the Boltzmann sigmoidal equation (n = 3).
